# Inorganic polyphosphate and the stringent response coordinately control cell division and cell morphology in *Escherichia coli*

**DOI:** 10.1101/2024.09.11.612536

**Authors:** Christopher W. Hamm, Michael J. Gray

## Abstract

Bacteria encounter numerous stressors in their constantly changing environments and have evolved many methods to deal with stressors quickly and effectively. One well known and broadly conserved stress response in bacteria is the stringent response, mediated by the alarmone (p)ppGpp. (p)ppGpp is produced in response to amino acid starvation and other nutrient limitations and stresses and regulates both the activity of proteins and expression of genes. *Escherichia coli* also makes inorganic polyphosphate (polyP), an ancient molecule evolutionary conserved across most bacteria and other cells, in response to a variety of stress conditions, including amino acid starvation. PolyP can act as an energy and phosphate storage pool, metal chelator, regulatory signal, and chaperone, among other functions. Here we report that *E. coli* lacking both (p)ppGpp and polyP have a complex phenotype indicating previously unknown overlapping roles for (p)ppGpp and polyP in regulating cell division, cell morphology, and metabolism. Disruption of either (p)ppGpp or polyP synthesis led to formation of filamentous cells, but simultaneous disruption of both pathways resulted in cells with heterogenous cell morphologies, including highly branched cells, severely mislocalized Z-rings, and cells containing substantial void spaces. These mutants also failed to grow when nutrients were limited, even when amino acids were added. These results provide new insights into the relationship between polyP synthesis and the stringent response in bacteria and point towards their having a joint role in controlling metabolism, cell division, and cell growth.

**IMPORTANCE:** Cell division is a fundamental biological process, and the mechanisms that control it in *Escherichia coli* have been the subject of intense research scrutiny for many decades. Similarly, both the (p)ppGpp-dependent stringent response and inorganic polyphosphate (polyP) synthesis are well-studied, evolutionarily ancient, and widely conserved pathways in diverse bacteria. Our results indicate that these systems, normally studied as stress-response mechanisms, play a coordinated and novel role in regulating cell division, morphology, and metabolism even under non-stress conditions.

## INTRODUCTION

Bacteria have evolved stress response systems to ensure their survival in everchanging environments and against host defenses. Two widely conserved stress response strategies are the general stress response in bacteria known as the stringent response, and the nearly universally conserved polyphosphate (polyP) pathway (1–4). However, the extent and nature of the interactions between these stress response pathways is poorly understood, as are their roles in bacterial physiology under non-stress conditions.

The stringent response is mediated by the small molecule guanosine 5’-diphosphate 3’-diphosphate (ppGpp) and guanosine 5’-triphosphate 3’-diphosphate (pppGpp), known collectively as (p)ppGpp (5–7). In *E. coli* (p)ppGpp is synthesized by the proteins RelA and SpoT (8–12), part of the RSH (**R**elA-**S**poT **h**omolog) family, with SpoT being able to both synthesize and degrade (p)ppGpp (8, 9, 13–17). SpoT is an essential gene in *relA*^+^ strains, as (p)ppGpp levels quickly rise to toxic levels in the absence of (p)ppGpp hydrolysis (18–22). (p)ppGpp is able to bind to and regulate the activity of proteins, including binding RNA polymerase and DksA to regulate genome-wide gene expression (23–25). (p)ppGpp downregulates DNA and RNA synthesis, protein production, and cell growth and modifies gene expression of up to one third of the genome in *E. coli* in response to a wide variety of stressors (5, 6, 14, 26–30). In recent years, (p)ppGpp has emerged as a master regulator of bacterial cellular biology and is now known to affect nearly every aspect from growth rate, sporulation, motility, competence, biofilm formation, toxin production and virulence of pathogenic bacteria (6, 27, 31–36). Relevant to the results we present here, the stringent response has also been linked to the downregulation of cell division, albeit by currently unknown mechanisms (37, 38).

Inorganic polyphosphate (polyP) is an evolutionarily ancient biopolymer and is found in nearly all bacterial and many eukaryotic cells (4, 39–43). In bacteria, polyP is involved in regulating gene expression, chelation of metals, acting as a protein-stabilizing chaperone, and can affect biofilm formation, stress sensing, quorum sensing, and motility (3, 4, 44–51). PolyP chains can be from dozens to thousands of phosphates long and are produced by polyphosphate kinase (PPK) and degraded by the enzyme exopolyphosphatase (PPX) (52–54). Pathogenic bacteria lose their virulence when polyP production is abolished, suggesting that polyP is vital for the ability of bacteria to cause harm, and motivating ongoing searches for PPK inhibitors as antivirulence drugs (4, 50, 55–58).

PolyP and the stringent response have long been suspected to be linked together, affecting virulence and the ability of bacteria to survive. Both *relA spoT* mutants lacking (p)ppGpp and *ppk* mutants lacking polyP have multiple amino acid auxotrophies and growth defects on minimal media, for example, suggesting parallel or linked roles in surviving nutritional stresses (47, 59, 60). (p)ppGpp is known to play a role in preventing degradation of polyP in bacteria by inhibiting PPX (42, 43, 61, 62), but evidence also suggests other potential links between these two fundamental systems (42, 63). Recent work from our lab showed that (p)ppGpp is not itself required for polyP synthesis, but did find a link between production of polyP in *E. coli* and the transcription factor DksA, which acts in coordination with (p)ppGpp to regulate gene expression within the cell (62, 64–66). Mutations of RNA polymerase that mimic the effects of (p)ppGpp binding to that enzyme on transcription (*a.k.a.* stringent alleles)(67–71) also reduce polyP synthesis by an unknown mechanism (42).

While the effects of (p)ppGpp and polyP during stress have been the focus of many investigations, their effect on cell physiology during normal growth conditions remains poorly understood, and the interaction between these two fundamental systems is not clear (6, 8, 13, 36, 42, 72). In this work, we identify striking and unexpected combinatorial phenotypes in *E. coli* mutants lacking both (p)ppGpp and polyP that suggest that these two pathways coordinately regulate fundamental mechanisms of cell division and morphology, even in the absence of nutritional stress. This provides new insights into the roles of general stress response pathways under non-stress conditions and raises important questions about the mechanisms bacteria use to maintain their integrity during growth, highlighting gaps in our knowledge in an area that has been the subject of many decades of research scrutiny.

## RESULTS

### Triple mutants lacking *ppk*, *relA*, and *spoT* have a growth defect on minimal medium that cannot be rescued with casamino acids

While both *ppk* and *relA spoT* mutants have well-described amino acid requirements in minimal media (43, 59, 73–75), both grow robustly on rich LB medium (**Fig 1**). We were surprised, therefore, to find that a triple *ppk relA spoT* mutant, entirely lacking the ability to synthesize both polyP and (p)ppGpp (73, 76), has a substantial growth defect on LB (**Fig 1**). Even more surprisingly, while we could easily rescue the growth defects of *ppk*, *relA*, *ppk relA*, and *relA spoT* mutants on minimal media by adding 0.05% (w/v) casamino acids (an undefined mixture of amino acids generated by acid hydrolysis of casein), casamino acids were unable to restore growth of the *ppk relA spoT* triple mutant on minimal medium. This indicated to us that there was an amino-acid independent defect in the *ppk relA spoT* strain that was not present in either parent strain, suggesting a previously unsuspected redundant metabolic role for polyP and (p)ppGpp in *E. coli*.

**FIG 1.**
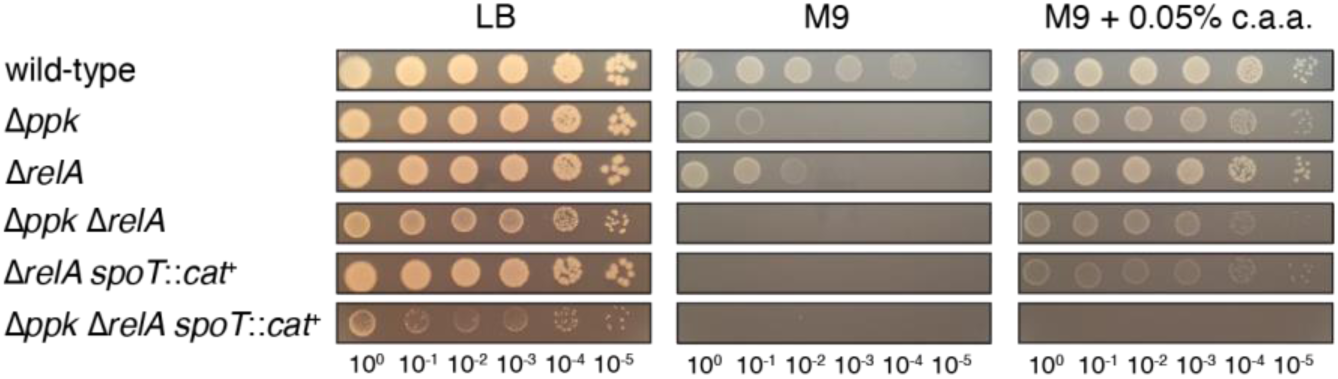
Triple mutants lacking *ppk*, *relA*, and *spoT* have a growth defect on minimal medium that cannot be rescued with casamino acids. *E. coli* strains MG1655 (wild-type), MJG0224 (MG1655 Δ*ppk-749*), MJG0226 (MG1655 Δ*relA782*), MJG1116 (MG1655 Δ*ppk-749* Δ*relA782*), MJG1136 (MG1655 Δ*relA782 spoT207*::*cat*^+^), and MJG1137 (MG1655 Δ*ppk-749* Δ*relA782 spoT207*::*cat*^+^) were grown overnight in LB broth, then rinsed and normalized to an A_600_ = 1 in PBS. Aliquots (5 µl) of serially-diluted suspensions were spotted on LB, M9 glucose, or M9 glucose containing 0.05% (w/v) casamino acids (c.a.a.) plates and incubated overnight at 37°C (representative image from at least 3 independent experiments).

*E. coli* lacking (p)ppGpp have low levels of the general stress response sigma factor RpoS (77, 78), and polyP has also been reported to increase RpoS expression (79), so we tested whether the phenotype of a *ppk relA spoT* mutant was also present in a *ppk rpoS* mutation, but it was not (**Supplemental Fig S1**) indicating that the role of (p)ppGpp in a *ppk* mutant is independent of RpoS-dependent transcriptional regulation (78). We also quantified guanosine nucleotide pools in *E. coli* in minimal media and found that (p)ppGpp accumulation was significantly higher in Δ*ppk* cells than in *ppk*^+^ strains (**Supplemental Fig S2**), supporting the idea that these two molecules can in some way compensate for each other’s absence.

### The growth defect of a *ppk relA spoT* mutant on minimal medium with casamino acids can be rescued by expression of either PPK or synthetase-active SpoT

Growth of the *ppk relA spoT* mutant on minimal medium containing casamino acids was restored by ectopic expression of either PPK or SpoT (**Fig 2A**). Complementation by SpoT was dependent on (p)ppGpp synthesis, since expression of a SpoT^D73N^ mutant allele lacking (p)ppGpp hydrolase activity (80) restored growth, but expression of SpoT^D259N^ and SpoT^D73N, D259N^ alleles, which lack (p)ppGpp synthetase activity (80), did not. Expression of wild-type SpoT enhanced the growth of a *relA spoT* mutant on minimal medium containing casamino acids (**Fig 2B**), but expression of SpoT alleles lacking (p)ppGpp synthetase activity did not, suggesting that (p)ppGpp synthesis underlies this effect. The *relA spoT* mutant could not be transformed with a plasmid encoding hydrolase-defective SpoT^D73N^ (data not shown). This was unsurprising, since accumulation of (p)ppGpp in *E. coli* cells lacking (p)ppGpp hydrolase activity is well known to prevent growth (73–75). The more surprising result from this experiment is that the *ppk relA spoT* mutant did tolerate SpoT^D73N^ expression (**Fig 2A**), suggesting that polyP is somehow involved in (p)ppGpp-dependent growth inhibition.

**FIG 2.**
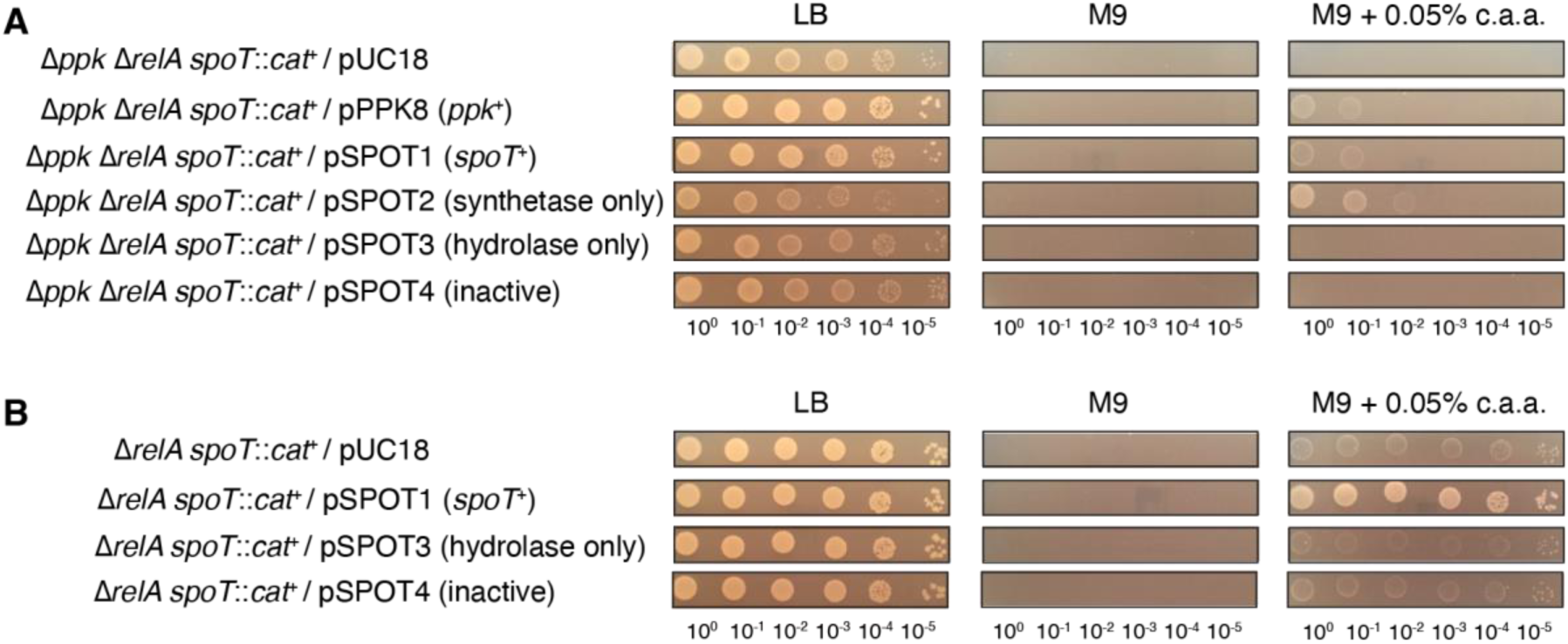
The growth defect of a *ppk relA spoT* mutant on minimal medium with casamino acids can be rescued by expression of either PPK or synthetase-active SpoT. *E. coli* strains (**A**) MJG1137 (MG1655 Δ*ppk-749* Δ*relA782 spoT207*::*cat*^+^) or (**B**) MJG1136 (MG1655 Δ*relA782 spoT207*::*cat*^+^) containing the indicated plasmids were grown overnight in LB broth containing ampicillin, then rinsed and normalized to an A_600_ = 1 in PBS. Aliquots (5 µl) of serially-diluted suspensions were spotted on LB, M9 glucose, or M9 glucose containing 0.05% (w/v) casamino acids (c.a.a.) plates containing ampicillin and incubated overnight at 37°C (representative image from at least 3 independent experiments).

### Both tryptone and yeast extract contain components that rescue the growth defect of a *ppk relA spoT* mutant on minimal medium with casamino acids

LB medium consists of 1% (w/v) of tryptone (a tryptic digest of casein), 0.5% (w/v) of yeast extract (prepared from the soluble fraction of boiled *Saccharomyces cerevisiae* cells), and 0.5% (w/v) of NaCl (81). Both tryptone and yeast extract at these concentrations restored growth of the *ppk relA spoT* triple mutant on minimal medium containing casamino acids (**Fig 3A**). The same effect could be seen in liquid media and with strains containing a complete deletion of *spoT* rather than the *spoT207*::*cat*^+^ insertion mutation (**Supplemental Fig S3**). Those experiments also showed that growth of all of the tested mutant strains in minimal media supplemented with a defined mixture of purified amino acids (yeast synthetic dropout mix supplement; Sigma Aldrich cat. #Y1501) was comparable to that in media supplemented with casamino acids, supporting our conclusions from **Fig 3A**. However, we found that variability was high in liquid media, especially later in the growth curves (possibly due to unpredictable accumulation of suppressor mutations), and opted to continue using solid media to evaluate the growth phenotype of the *ppk relA spoT* mutant.

**FIG 3.**
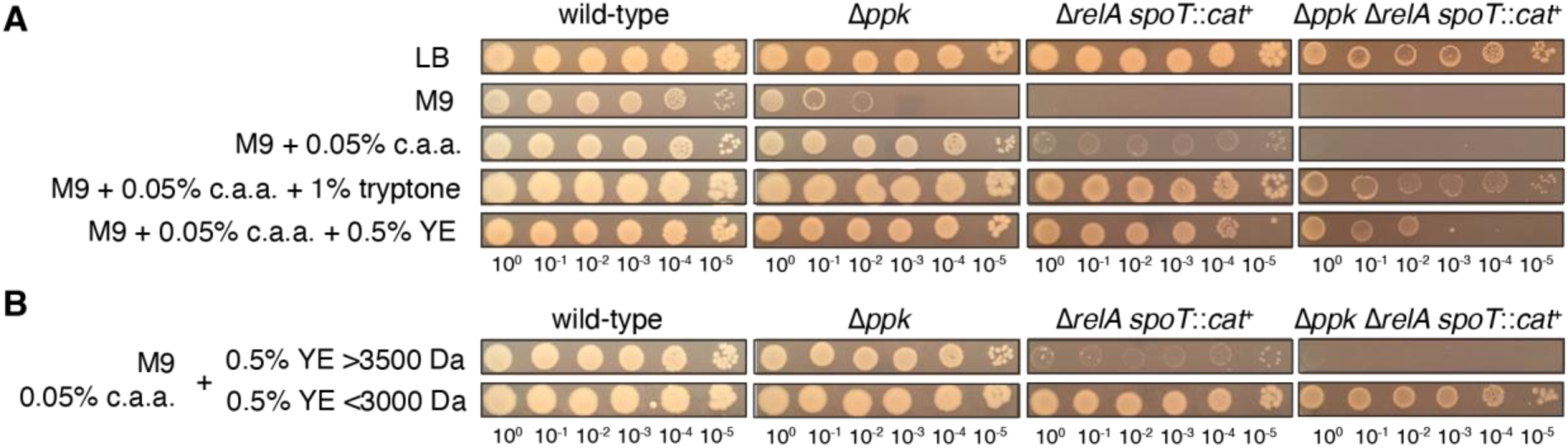
Both tryptone and yeast extract contain components that rescue the growth defect of a *ppk relA spoT* mutant on minimal medium with casamino acids. *E. coli* strains MG1655 (wild-type), MJG0224 (MG1655 Δ*ppk-749*), MJG1136 (MG1655 Δ*relA782 spoT207*::*cat*^+^), and MJG1137 (MG1655 Δ*ppk-749* Δ*relA782 spoT207*::*cat*^+^) were grown overnight in LB broth, then rinsed and normalized to an A_600_ = 1 in PBS. Aliquots (5 µl) of serially-diluted suspensions were spotted on LB, M9 glucose, or M9 glucose containing the indicated percentages (w/v) of casamino acids (c.a.a.) and (**A**) tryptone or yeast extract (YE) or (**B**) yeast extract fractions containing compounds either greater than 3500 Da or less than 3000 Da and incubated overnight at 37°C (representative image from at least 3 independent experiments).

We used filtration and dialysis to divide yeast extract into fractions with molecular weights greater than 3500 Da or less than 3000 Da and found that the component(s) responsible for restoring growth of the *ppk relA spoT* mutant on minimal medium containing casamino acids were present in the small molecule (*e.g.* less than 3000 Da) fraction (**Fig 3B**). Because LB medium is regularly sterilized by autoclaving, the compound(s) in question must also be heat-stable, and we are currently working to identify the relevant molecule(s), which we expect to give insights into the metabolic pathway(s) responsible for the inability of the *ppk relA spoT* mutant to grow on minimal media.

### Morphological defects of *ppk relA spoT* cells 1: filamentation

Since the *ppk relA spoT* mutant did not grow as well on LB plates as the other mutant strains, we wanted to see if there were visible morphological defects present in this strain or its parent strains. We performed confocal time-lapse microscopy and noted that cells lacking either polyP or (p)ppGpp grown in LB are longer than wild-type cells, indicating defects in cell division (**Fig 4**). Our results showed that a *ppk* mutant lacking polyP was slightly longer than the wild type MG1655 while growing in exponential phase on LB (**Fig 4**). Strains deficient in (p)ppGpp were also filamentous, with a *relA* mutant being longer on average than a double *relA spoT* mutant (**Fig 4**). The *ppk relA* mutant lacking polyP and only containing SpoT for production of (p)ppGpp were about the same length as the *relA* mutant or the *relA spoT* mutant (**Fig 4**). These results are in general agreement with previous work in *E. coli* showing increased cell length in *relA spoT* mutants (82) and in *Pseudomonas aeruginosa* showing that strains lacking polyP become filamentous in stationary phase (83). The triple mutant *ppk relA spoT* contained filamentous cells as well (**Fig 4**), however these cells were much more heterogenous than any of the other mutants, appearing less stable with odd morphologies that we will explore in depth in the following sections. When grown on MOPS minimal media agarose pads, the various mutant cells all had morphologies similar to the wild type and were no longer filamentous (**Supplemental videos SV1** and **SV2**). The *ppk relA spoT* triple mutant immediately stopped growing on the MOPS agarose pad and slowly started to shrink over several hours (**Supplemental video SV3**).

**FIG 4.**
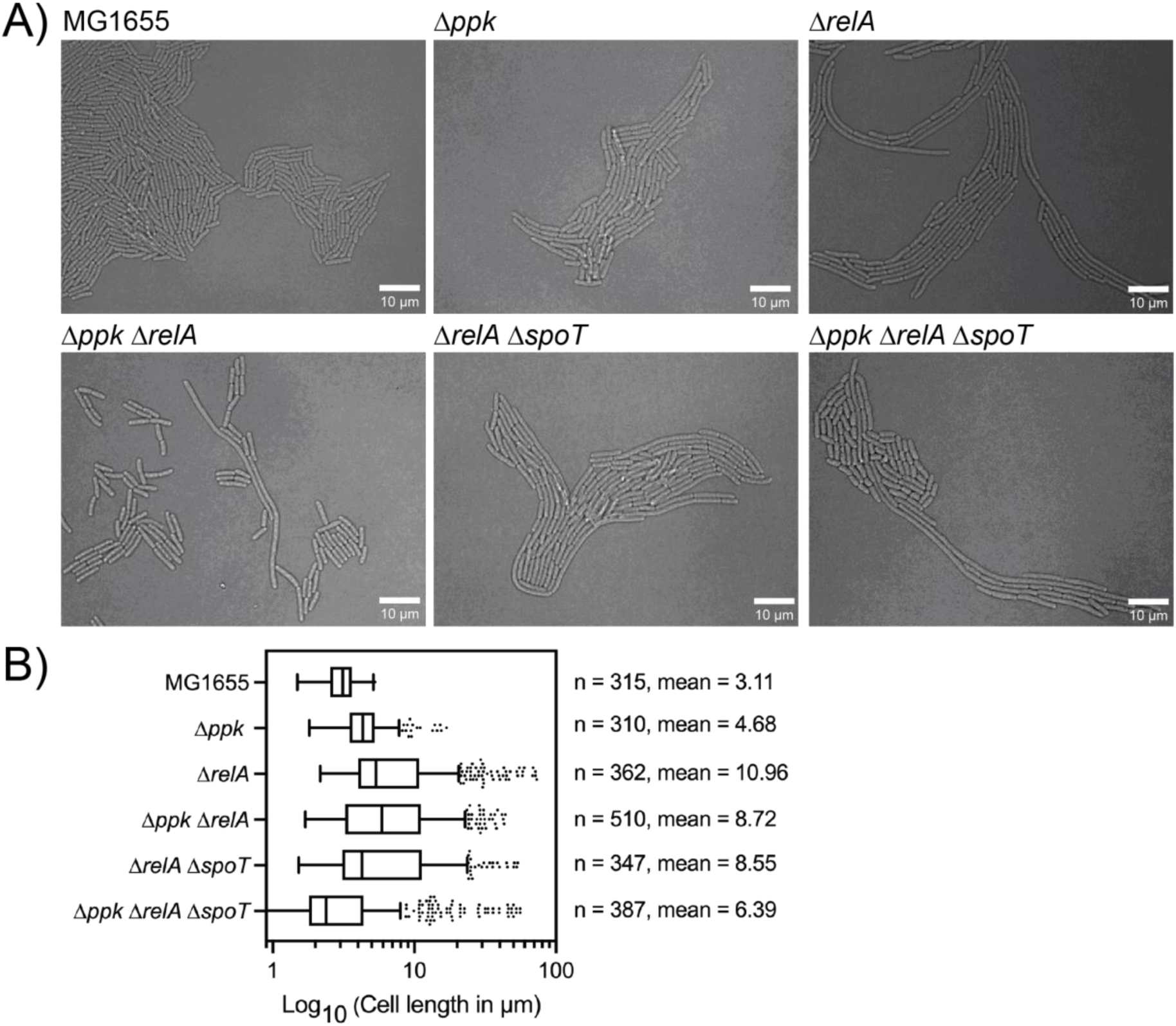
Microscopy of polyP and (p)ppGpp mutants shows filamentous cell growth. (**A**) Confocal microscopy images of MG1655 (MJG0001), *Δppk* (MJG0224), *ΔrelA* (MJG0226), *Δppk ΔrelA* (MJG1116), *ΔrelA ΔspoT* (MJG1287), and *Δppk ΔrelA ΔspoT* (MJG1282). All images were captured on LB agarose pads incubated at 37°C and imaged every 5 minutes while growing. Length of cells was determined and manually calculated using FIJI. (**B**) Cell lengths were calculated from images captured and analyzed in FIJI. Data calculated in Prism GraphPad, all mutant population averages are significantly different from MG1655 with a P value <0.0001 (calculated by one-way ANOVA in Prism) except for MJG0224 (Δ*ppk*).

### Morphological defects of *ppk relA spoT* cells 2: displaced FtsZ ring formation and deviant cell division

To determine where cell division was taking place in these filamentous cells, we used a FtsZ-GFP reporter to observe where the Z-ring was localizing within cells during cell division (84). During time-lapse microscopy, we observed the FtsZ reporter forming a Z-ring at the midpoint of wild-type cells as expected (**Supplemental Fig S4** and **Supplemental Video SV4**) (85–87). The triple mutant *ppk relA spoT* however showed that, while some FtsZ rings formed normally, there were many cells with profoundly disrupted Z-ring localization phenotypes. We observed cells containing multiple Z-rings within a single cell (**Fig 5A and 5B**), including instances with two Z-rings forming in the middle of a cell with both Z-rings constricting and resulting in the release of a non-growing mini-cell (**Fig 5A**).

**FIG 5.**
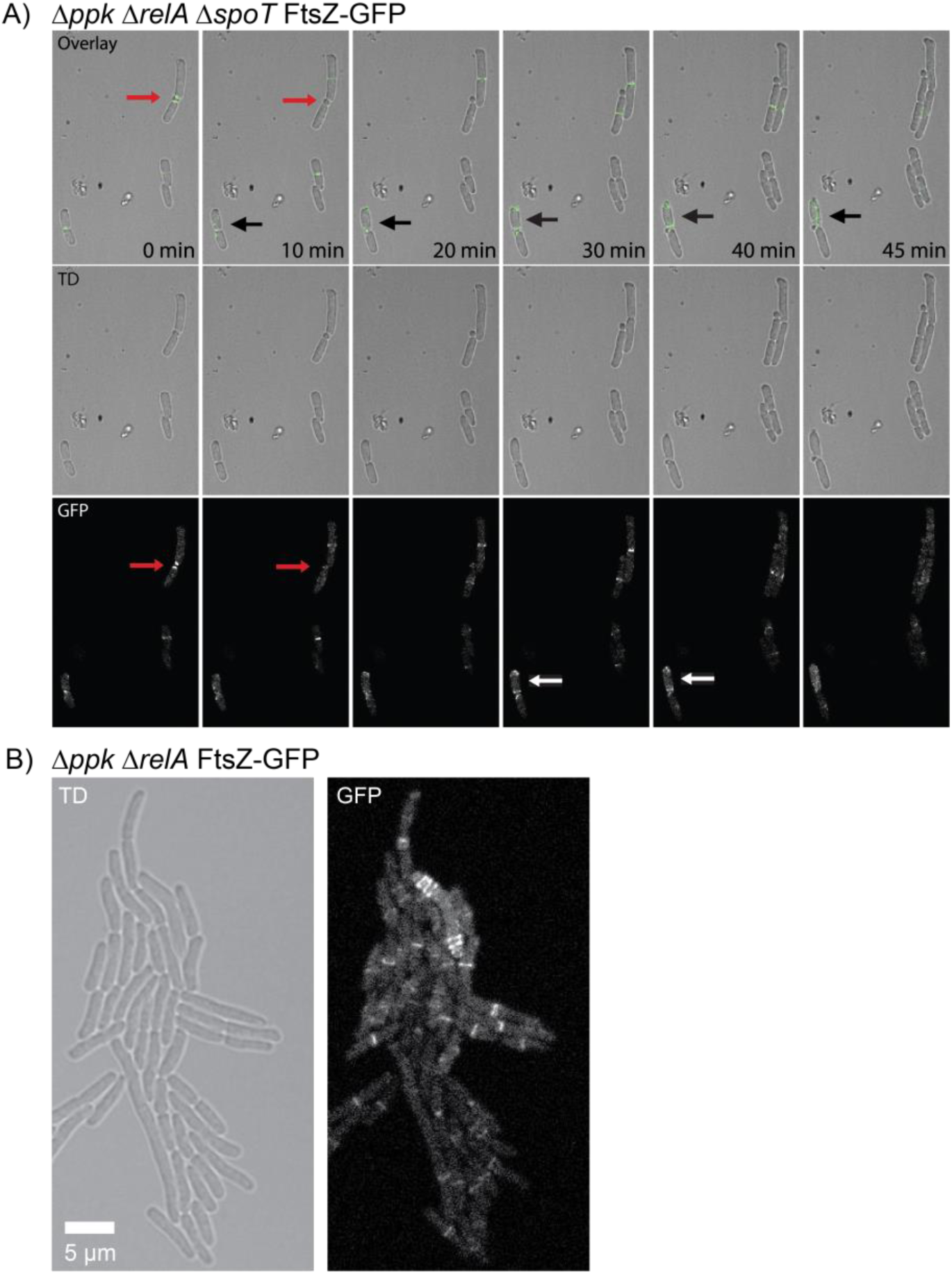
Strains lacking polyP and (p)ppGpp have disrupted cell division. (**A**) Confocal fluorescence time-lapse microscopy of the mutant *ppk relA spoT* FtsZ-GFP (MJG2405) on an LB agarose pad at 37°C. The triple mutant forms two Z-rings in the middle of the cell, releasing a mini cell (red arrows). There are also two Z-rings which form at either pole of a single cell, both functional and releasing a mini cell (white arrows). Example of mini-cell formation and release can be seen in CEM (S4). (**B**) Confocal fluorescence time-lapse microscopy of the mutant *ppk relA* FtsZ-GFP (MJG2403) on an LB agarose pad at 37°C. This image shows a mutant forming three Z-rings at one pole, and at least 3 at the opposite pole as well, with no Z-rings forming in the middle of the cell, for a total of six Z-rings in a single cell. This cell continued to grow without lysing (**Supplemental Video SV5**).

Not only were multiple Z-rings forming at the midpoint of cells, but we also observed Z-rings forming at the poles of cells dividing and releasing mini-cells (**Figs 5A,B**, **Supplemental Video SV5**). We also observed an FtsZ ring forming at the side wall of a branching cell, pinching off and releasing a mini-cell from the side wall of a branching cell (**Supplemental Fig S5**). Most cells did appear to divide normally when cell division occurred, with filamentous cells failing to localize a Z-ring until division occurred. The *ppk* and *relA spoT* mutants did not have these errors with FtsZ formation and formed Z-rings normally in LB at 37°C (**Supplemental Fig S4**).

When the *ppk relA spoT* mutant was observed by transmission electron microscopy (TEM), we confirmed these observations (**Supplemental Figs S6** and **S7**) and noticed other cell division defects. We observed cells appearing to divide, but without forming a proper septum fully separating the dividing cells (**Fig 6**). We observed a cell where cell division was apparently aborted, as while division had mostly occurred and the cytoplasm was divided, the two cells were still connected by a membrane bridge (**Fig 6A** and **6B**). We also observed cells where cell division was actively occurring, however with a bridge between both cells still connecting the cytoplasm from one cell to another (**Fig 6C** and **6D**). These cells appear to have aborted cell division at different times, suggesting unstable FtsZ ring formation, where FtsZ did not complete cell division and septum formation. These results suggest an instability in FtsZ ring formation when cells lack both polyP and (p)ppGpp, connoting a link between polyP and (p)ppGpp redundantly stabilizing FtsZ ring formation during cell division.

**FIG 6.**
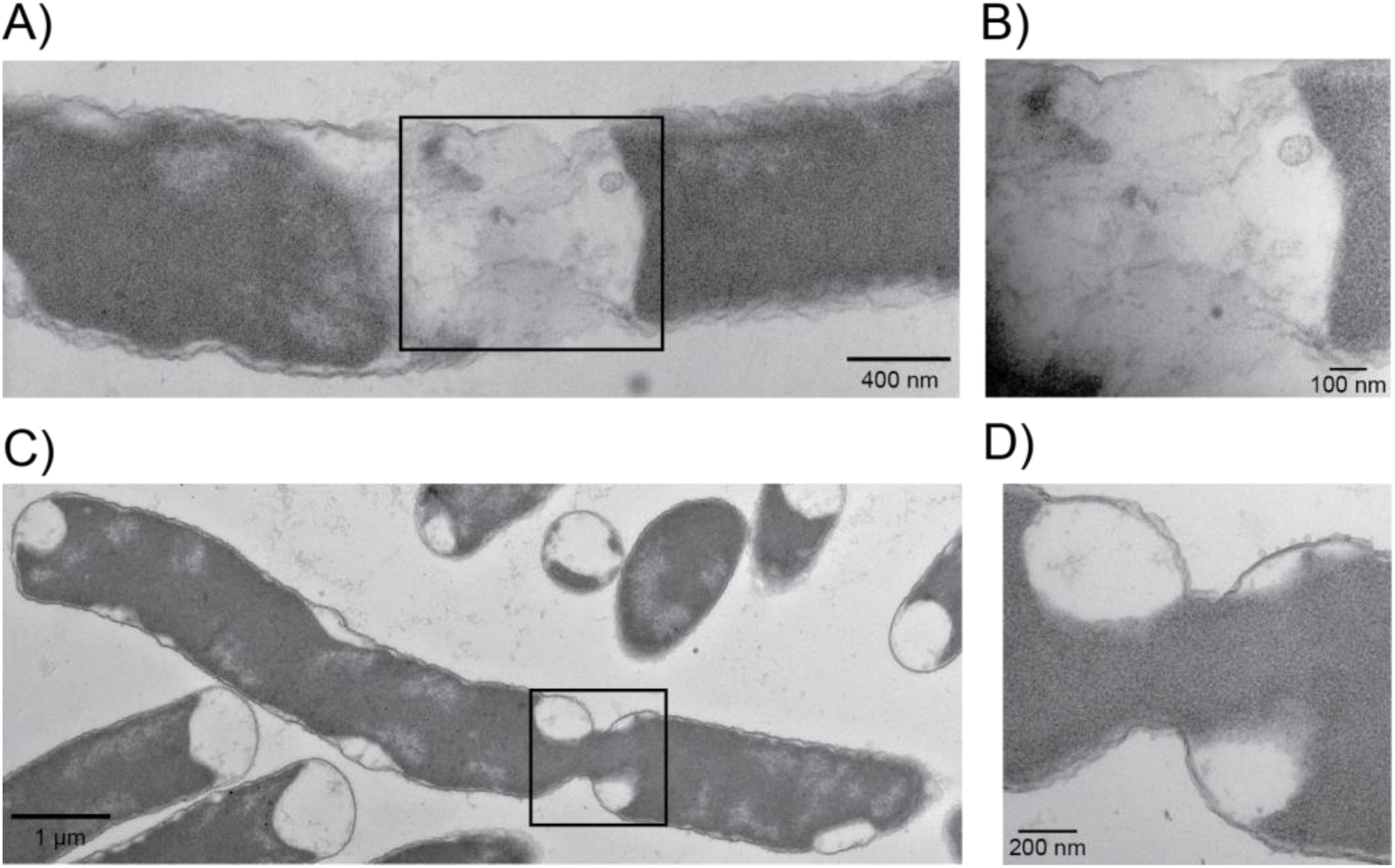
Transmission Electron Microscopy (TEM) of *ppk relA spoT* cells failing to divide properly. **A)** TEM of *ppk relA spoT* (MJG1282) failing to divide properly. This cell cytoplasm has condensed away from the divisisome site but is still connected by a small bridge and could be the result of disrupted FtsZ-ring formation. **B)** This image is the square section from **A)** at a higher magnification. **C)** TEM of *ppk relA spoT* (MJG1282) not completely forming a divisisome and staying connected through a bridge while still sharing cytoplasmic contents. **D)** This image is the square section from **C)** at a higher magnification.

### Morphological defects of *ppk relA spoT* cells 3: branched cells

While observing the *ppk relA* double mutant and the *ppk relA spoT* triple mutants we discovered that these mutants were able to form very unusual branched cells, along with other odd and unexpected cell morphologies. Only cells lacking both polyP and (p)ppGpp were able to form branched cells, including instances of cells with more than three distinct poles (**Fig 7A-C**, **Supplemental Fig S5**). In the *ppk relA* double mutant we were able to observe a branched cell developing over time, with FtsZ still capable of forming Z-rings and causing cell division (**Fig 7A**). The double mutant *ppk relA* developed branched cells less frequently (∼2.4% of cells) than the triple mutant *ppk relA spoT* (∼3.8% of cells), but there was less uniformity in branched cells of the triple mutant. The triple mutant developed into more heterogenous branching cells with more noticeable defects in cell wall morphology including formation of spheroplasts (**Fig 7C** and **Supplemental Videos SV6**-**8**). The development of branched cells did not depend on the presence of the FtsZ-GFP reporter, as we observed the same morphologies in the strains lacking the reporter (**Fig 7C**). These results suggest a combined or redundant role for polyP and (p)ppGpp in regulating cell wall synthesis and/or integration of newly synthesized peptidoglycan.

**FIG 7.**
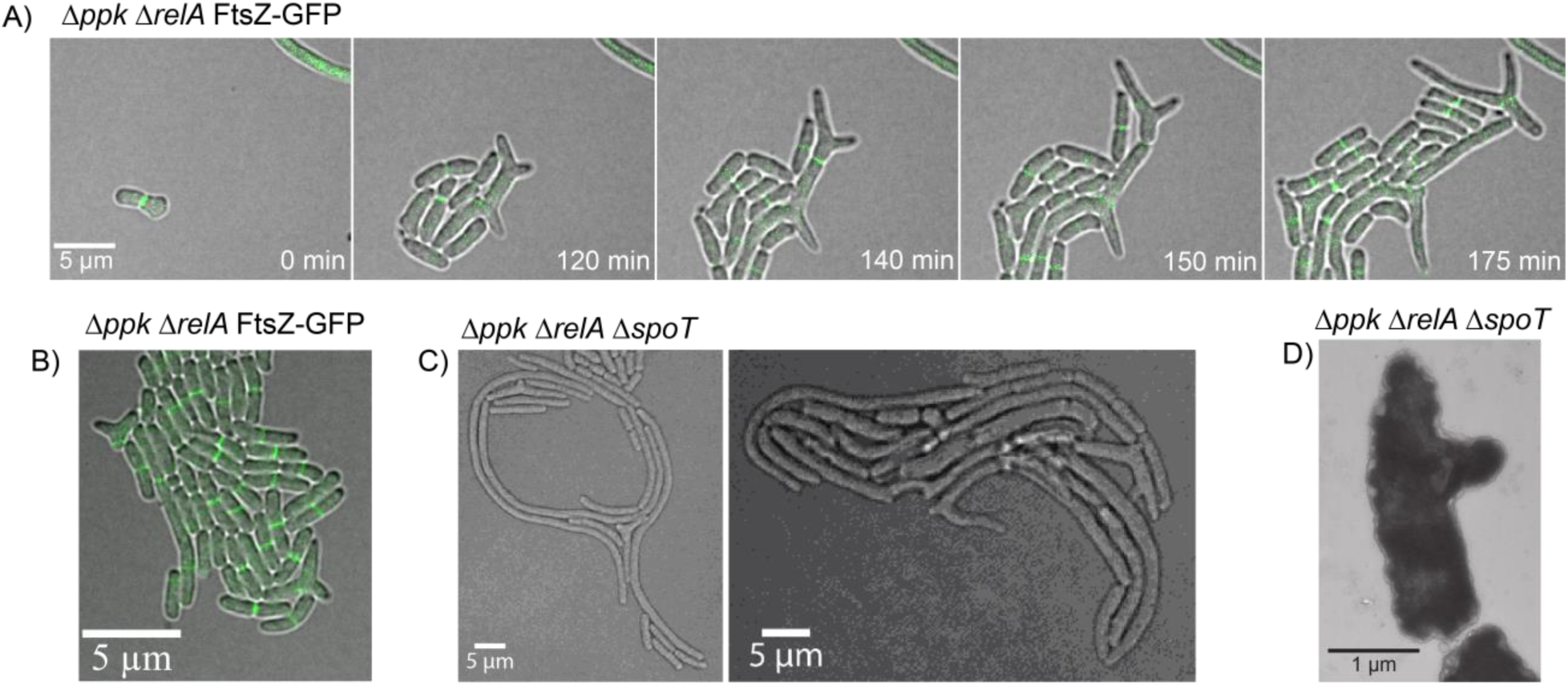
Cells lacking polyP and (p)ppGpp can develop branching cell morphologies. (**A**) Confocal fluorescence time-lapse microscopy of the mutant *ppk relA* FtsZ-GFP (MJG2403) on an LB agarose pad at 37°C showing branching cells. Branched cells are still capable of dividing and growing. (**B**) Still image of confocal fluorescent microscopy of *ppk relA* FtsZ-GFP mutant (MJG2403) mutant on LB agarose pad at 37°C showcasing branched cells. (**C**) Still image of confocal microscopy of *Δppk ΔrelA ΔspoT* (MJG1282) mutant on LB agarose pad at 37°C showcasing branched cells and other very odd cell morphologies. (**D**) Transmission Electron Microscopy of *ppk relA spoT* (MJG1282) grown in LB at 37°C until log phase prior to imaging. This image shows a cell developing a branch point during growth.

### Morphological defects of *ppk relA spoT* cells 4: cell envelope defects and cytoplasmic condensation

Another unexpected phenotype we noted while observing the *ppk relA spoT* triple mutant was the presence of cells developing what appear as “holes” or void spaces within the bacterial cell, appearance of which preceded the leakage of cytoplasmic contents out of the cell, as represented by the FtsZ-GFP fusion protein (**Fig 8A**). To investigate this phenotype more closely, we performed cryo-electron microscopy of the triple mutant, looking for evidence of disrupted cellular membranes. We imaged what appears to be an invagination of the cell wall, with cytoplasmic contents being released as blebs from the cell surrounded by a cell wall and membranes (**Fig 8B**). The site where cytoplasmic contents appear to be “leaking” from may be a site of cell division, as we saw similar invaginated structures form in dividing cells (**Supplemental Figs S8**, **S9**). We also observed disruptions in the outer membrane of the triple mutant (**Fig 8B**), with what appears to be cytoplasmic contents condensing oddly within the cell (**Fig 8B** and **Supplemental Figs S10-13**).

**FIG 8.**
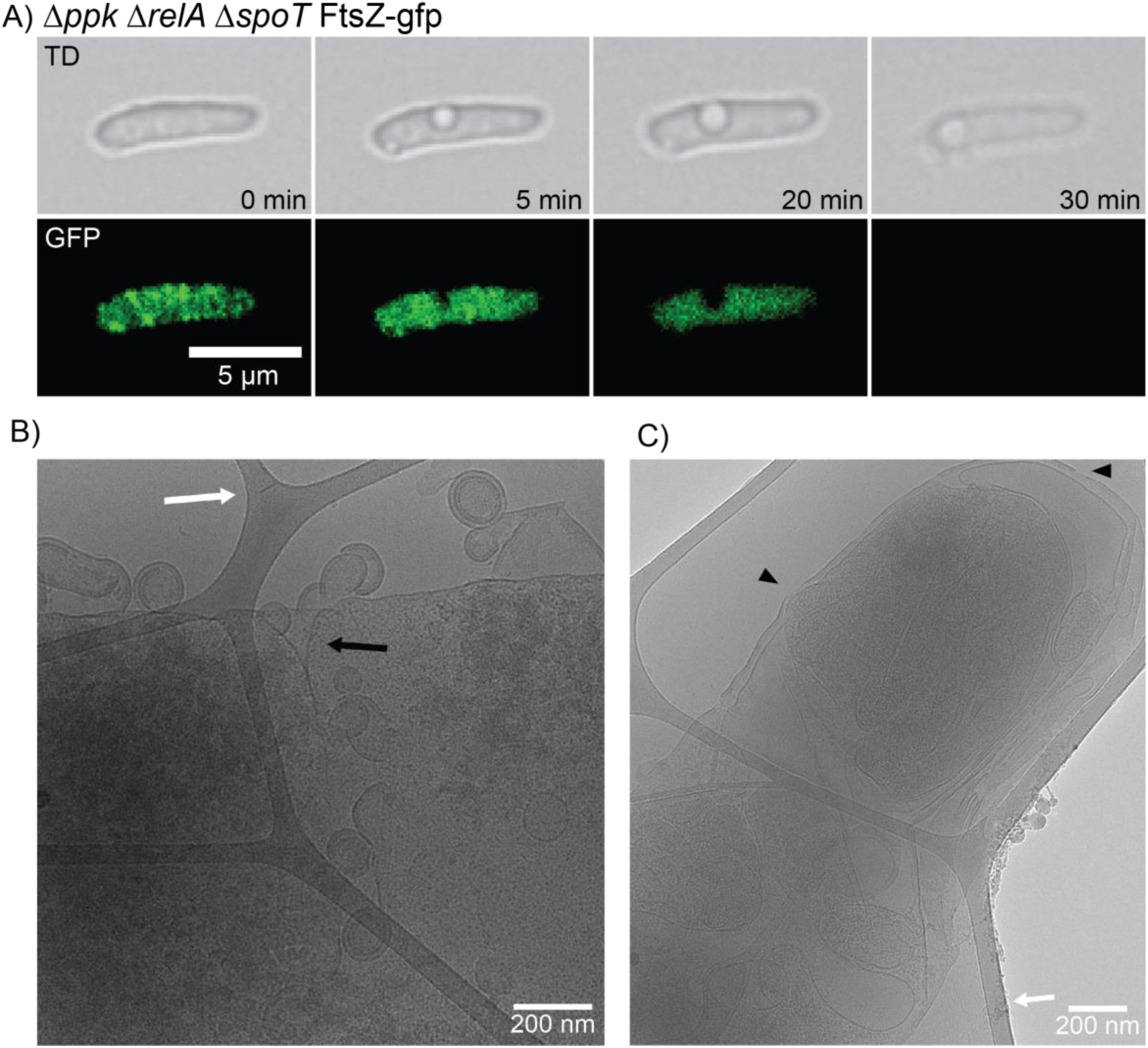
*ppk relA spoT* triple mutant cells can develop perforated membranes, leaking cytoplasmic contents out of the cell. (**A**) Time-lapse fluorescent microscopy of the *ppk relA spoT* mutant with the FtsZ-GFP reporter (MJG2405) showing empty space within the cell. FtsZ appears to fail to localize within the cell prior to losing its cytoplasmic contents just before cell death occurs. (**B**) CEM image of *ppk relA spoT* mutant (MJG2405) showing what appears to be a leaking cell wall with cytoplasmic contents blebbing off. (**C**) CEM image of *ppk relA spoT* mutant (MJG2405) showing what appears to be holes in the cellular membrane and cytoplasm condensation such as from a loss of liquid from within the cell.

Using TEM, we observed *ppk relA spoT* mutant cells that appeared to have their cytoplasm and inner membrane shrunk away from the cell wall, creating large periplasmic spaces, a phenomenon known as plasmolysis (**Figs 8** and **9**, **Supplemental Figs S10** and **S11**). Plasmolysis has been known to be caused by hyperosmotic shock in bacteria, although in that case the cytoplasmic shrinkage is relatively uniform among cells in a population (88, 89). This suggests a possible mechanism for polyP and (p)ppGpp to co-regulate response to osmotic stress, or potential for dysregulation of efflux pumps causing the loss of cytoplasmic contents and subsequent efflux of water. This loss of water, however, could also be due to loss of water through a leaking membrane (**Fig 8**) or other unknown mechanisms. In another paper reporting similar-appearing void spaces in bacterial cytoplasms, the authors believed they were seeing cytoplasmic condensation caused by disruptions in the cell envelope due to treatment with sublethal concentrations of antibiotics (90), in which case our observations might imply a role for polyP and (p)ppGpp in synthesis and/or incorporation of the newly synthesized peptidoglycan into the cell wall. Regardless of the underlying mechanism(s), these results indicate that in a *ppk relA spoT* mutant, some cells are unable to properly maintain their growing cell wall or membranes.

**FIG 9.**
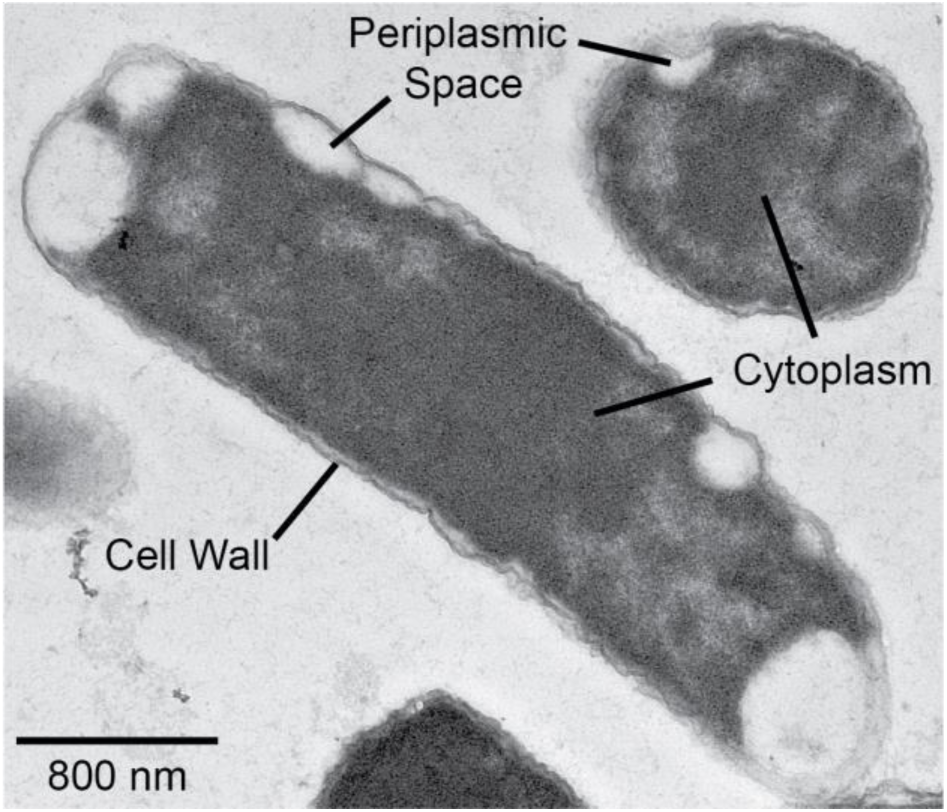
TEM of *ppk relA spoT* mutant showing plasmolysis in both cross-sectional and trans-sectional viewpoint. TEM of *ppk relA spoT* (MJG1282) showing the inner membrane appearing to shrink and pull away from the cell wall (plasmolysis), leaving large periplasmic spaces within the cell.

### Stringent alleles of RNA polymerase restore some, but not all phenotypes of a *ppk relA spoT* mutant

The amino acid auxotrophy of *relA spoT* mutants can be rescued by mutations in RNA polymerase called stringent alleles, which mimic the regulatory effect of (p)ppGpp binding to RNA polymerase on transcription, including activation of expression of amino acid synthesis operons (68, 69, 74, 75). Many of these mutations in RNA polymerase also confer rifampicin resistance (91). The stringent alleles *rpoB3443* and *rpoB3449* (69) were, as expected, able to restore growth of a *relA spoT* mutant in the absence of casamino acids, while the rifampicin-resistant but non-stringent allele *rpoB148* (91) did not (**Fig 10A**). Notably, *rpoB3443* and *rpoB3449* restored growth of the *ppk relA spoT* triple mutant in the presence of casamino acids, but not in their absence. Microscopic observation of a *ppk relA spoT rpoB3443* mutant grown on LB, however, revealed a dramatic restoration of wild-type cell morphology (**Fig 10B** and **Supplemental Video SV9**). The growth rates of the *ppk relA spoT* and *ppk relA spoT rpoB3443* mutants in LB, while slower than wild-type, were not different from each other (**Supplemental Fig S14**), so this morphological rescue was not due to a change in growth rate.

**FIG 10.**
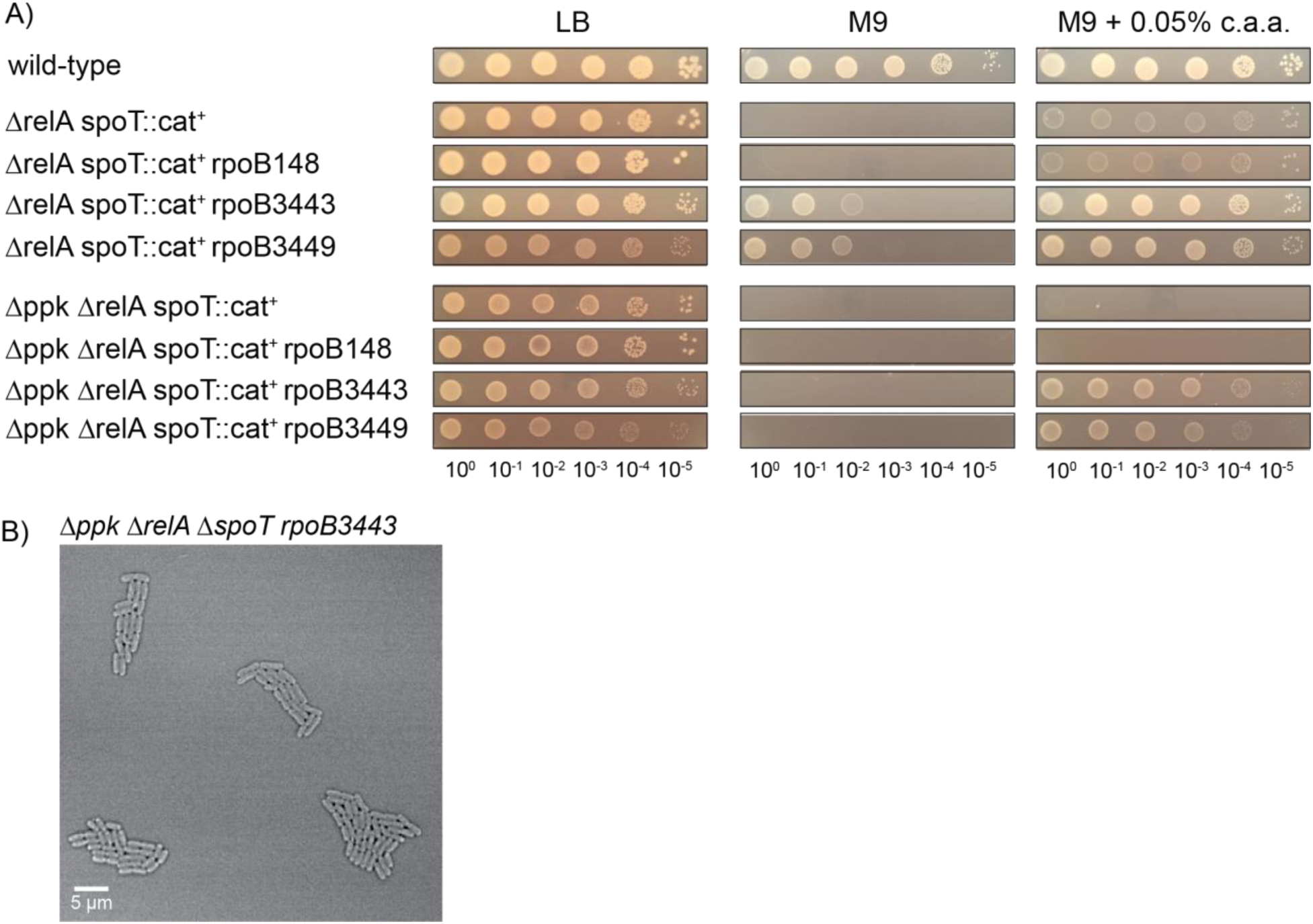
Stringent alleles of RNA polymerase restore growth of *ppk relA spoT* mutants on minimal medium with casamino acids. (**A**) *E. coli* strains MG1655 (wild-type), MJG1136 (MG1655 Δ*relA782 spoT207*::*cat*^+^), MJG1137 (MG1655 Δ*ppk-749* Δ*relA782 spoT207*::*cat*^+^), MJG1237 (MG1655 Δ*relA782 spoT207*::*cat*^+^ *rpoB3449*), MJG1241 (MG1655 Δ*ppk-749* Δ*relA782 spoT207*::*cat*^+^ *rpoB3449*), MJG1579 (MG1655 Δ*relA782 spoT207*::*cat*^+^ *rpoB3443*), MJG1580 (MG1655 Δ*relA782 spoT207*::*cat*^+^ *rpoB148*), MJG1581 (MG1655 Δ*ppk-749* Δ*relA782 spoT207*::*cat*^+^ *rpoB3443*), and MJG1582 (MG1655 Δ*ppk-749* Δ*relA782 spoT207*::*cat*^+^ *rpoB148*) were grown overnight in LB broth, then rinsed and normalized to an A_600_ = 1 in PBS. Aliquots (5 µl) of serially-diluted suspensions were spotted on LB, M9 glucose, or M9 glucose containing 0.05% (w/v) casamino acids (c.a.a.) plates and incubated overnight at 37°C (representative image from at least 3 independent experiments). (**B**) Confocal microscopy of *ppk relA spoT rpoB3443* (MJG1581) on a LB agarose pad at 37°C.

These results taken together suggest that the nutritional (**Figs 1**-**3** and **10A**) and morphological (**Figs 4-9** and **10B**) phenotypes of the *ppk relA spoT* mutant can be genetically separated and that the morphological phenotypes of this strain in particular appear to be linked to transcriptional regulation by (p)ppGpp. In contrast, the combinatorial growth defect of the triple mutant on minimal medium may be dependent on (p)ppGpp’s impacts on a protein or proteins other than RNA polymerase (65, 74, 75). These results also reinforce our conclusion that whatever compound(s) are present in LB that allow growth of the triple mutant (**Fig 3**) are not likely to be amino acids.

## DISCUSSION

Connections between (p)ppGpp and polyP in *E. coli* have been suspected for decades (43, 61), but the nature and consequences of those connections have remained obscure (41, 42, 46, 92). We have now identified a striking combinatorial phenotype which clearly demonstrates that these two conserved “stress response” molecules play important linked roles in controlling fundamental metabolic processes under non-stress growth conditions. The fact that these phenotypes appear only when both (p)ppGpp and polyP are eliminated suggests that either one alone is sufficient to maintain more or less normal cells, and that therefore some critical pathway or pathways must be regulated by both molecules. The challenge that remains is to dissect the mechanism(s) by which (p)ppGpp and polyP coordinate these processes.

Both (p)ppGpp and polyP can act at multiple regulatory levels in the bacterial cell. The regulatory consequences of (p)ppGpp synthesis are better studied and include dramatic changes in the genome-wide transcriptome of *E. coli* due to (p)ppGpp binding to RNA polymerase and the transcription factor DksA (2, 62). (p)ppGpp can also direct regulation of a variety of other enzymes (1, 26, 65), including notably the exopolyphosphatase PPX, which is inhibited by (p)ppGpp (43, 54, 61). More than 700 genes are transcriptionally regulated by (p)ppGpp, approximately 400 of which are inhibited and 300 are stimulated (including, to a modest extent, *ppk*)(93), but the exact list depends on growth conditions and on how (p)ppGpp synthesis is induced (93, 94). Although stringent alleles of RNA polymerase have been known for many decades and are thought to mimic the effects of (p)ppGpp binding (68, 69, 74, 75), to our knowledge no genome-wide characterization of their impact on transcription has been performed.

More than 50 other proteins that bind (p)ppGpp in *E. coli* have been identified (65), and the impact of that binding has been characterized for only a subset of those proteins (1, 26, 65).

PolyP can also act at multiple levels, although its impact is considerably less well characterized. PolyP interacts directly with the Lon protease, for example, to modulate its activity and substrate specificity (95–98), which could affect a very broad range of potential protein targets in the cell both directly and indirectly. In combination with Hfq, polyP is also able to silence transcription of some genes (99), and a Δ*ppk* mutant has substantial changes in its transcriptome and proteome (100), although how much of this is due to direct regulatory impacts as opposed to indirect responses to a lack of polyP remains unclear. Whether other *E. coli* proteins might be impacted by polyP binding is not well known, since to our knowledge no systematic study identifying polyP-binding proteins in bacteria has been performed (101, 102). The fact that we can genetically distinguish between the metabolic and morphological phenotypes of *ppk relA spoT* mutants (**Fig 10**) strongly suggests that at least two different pathways are co-regulated by (p)ppGpp and polyP, which further complicates the problem.

Based on the existing literature, we can identify a few potential overlaps between the (p)ppGpp and polyP regulons in *E. coli* which might be relevant to the phenotypes we observe in *ppk relA spoT* mutants. As one example, (p)ppGpp binds to the DapB protein (65) and upregulates transcription of many of the *dap* genes involved in the synthesis of the peptidoglycan precursor diaminopimelate (93, 103), while in a Δ*ppk* mutant, Varas *et al.* (100) reported an increase in DapA protein levels and a decrease in *dapF* transcription. In another example, FtsY, an essential component of the signal recognition particle that delivers integral membrane proteins to the inner membrane (104), is allosterically inhibited by (p)ppGpp binding (65, 105), modestly downregulated transcriptionally by (p)ppGpp (93), and its transcription is increased in a Δ*ppk* mutant (100). Either of these pathways could conceivably contribute to the cell envelope defects we observe in *ppk relA spoT* mutants (**Figs 7**-**9**), but since each of those experiments was performed under different conditions and in varying strain backgrounds it is not possible to make definite conclusions about whether these potential points of regulatory overlap are either real or meaningful, and certainly not about whether they contribute to the phenotypes of *ppk relA spoT* strains. We are currently working to characterize the transcriptional and post-transcriptional impacts of (p)ppGpp and polyP on *E. coli* under growth conditions where we observe *ppk relA spoT* phenotypes, with the goal of systematically identifying genes, proteins, and pathways impacted by both molecules.

Perhaps the most surprising and most difficult to explain of the phenotypes of the *ppk relA spoT* mutant is the mislocalization of Z-rings (**Figs 5**-**7**). The positioning of FtsZ at the center of the cell is a highly regulated part of cell division (106). *E. coli* has two major systems for preventing formation of Z-rings anywhere other than mid-cell: the MinCDE system that inhibits Z-ring formation at the poles (107) and the SlmA nucleoid exclusion protein that prevents Z-ring formation around the DNA nucleoid (108). Neither of these systems has been reported to be impacted by either polyP or (p)ppGpp, although examination of published transcriptome data indicates that *minD* and *minE* are slightly upregulated and *slmA* is slightly downregulated by (p)ppGpp (93). Overexpression of FtsZ can result in production of multiple Z-rings within cells, including both mid-cell and polar localizations (109), reminiscent of some of our observations of *ppk relA spoT* mutants containing many concurrent Z-rings (**Fig 5**). FtsZ regulation is complex, but not known to depend on either (p)ppGpp or polyP (106). Our data strongly suggest that there are important unknowns remaining in our understanding of the regulation of proper cell division in *E. coli*.

In *E. coli*, both (p)ppGpp and polyP are present at very low levels under non-stress conditions and those levels are strongly increased by various stresses, which is critical for the ability of the bacteria to survive those stress treatments (2–4). However, the phenotypes we report here for the *ppk relA spoT* mutant are in rich LB medium, meaning that those phenotypes depend on unstimulated basal levels of (p)ppGpp and polyP. There is an increasing appreciation in the literature that basal levels of (p)ppGpp make important contributions to diverse aspects of *E. coli* physiology (110), including cell division (111). The same is likely to be true of basal polyP levels, since substantial changes in gene expression and proteome composition are seen in Δ*ppk* mutants grown in rich medium (99, 100).

A final point worth considering is the surprising heterogeneity of the *ppk relA spoT* cells. As shown in **Figs 4**-**9**, not every cell of the triple mutant has the same morphological defects. Some of them look fairly normal, some are filamentous, some have mislocalized Z-rings, and a few develop more severe defects, including branching, spheroplast formation, void spaces, or lysis. What underlies this diversity and what distinguishes individual cells that do well from cells that do not is an intriguing unanswered question.

## MATERIALS AND METHODS

### Databases and primer design

We obtained gene and protein sequences and other information from the Integrated Microbial Genomes database (112) and from EcoCyc (103), and designed PCR and sequencing primers with Web Primer (www.candidagenome.org/cgi-bin/compute/web-primer). Mutagenic primers were designed with PrimerX (www.bioinformatics.org/primerx/index.htm).

### Bacterial strains and growth conditions

All strains used in this study are listed in **Table 1**. We grew *E. coli* at 37°C in Lysogeny Broth (LB)(113) containing 5 g l^-1^ NaCl, in M9 minimal medium (81) containing 4 g l^-1^ glucose, or in MOPS minimal medium (114) containing 2 or 4 g l^-1^ glucose. Solid media contained 1.5% (w/v) agar (Becton Dickinson cat. #214010). Ampicillin (100 µg ml^-1^), chloramphenicol (17.5 or 35 µg ml^-1^), kanamycin (25 or 50 µg ml^-1^), or rifampicin (50 µg ml^-1^) were added when appropriate. Nutritional supplements used were yeast synthetic dropout mix supplement (Sigma Aldrich cat. #Y1501), casamino acids (Fisher Scientific cat. #BP1424), tryptone (Fisher Scientific cat. #BP1421), or yeast extract (Becton Dickinson cat. #288620). For growth curves, *E. coli* strains of interest were grown overnight at 37°C with shaking in LB, then normalized to a A_600_ = 1 and rinsed three times with sterile PBS. The resulting cell suspensions were diluted 1:40 into fresh media. Growth curves were performed in clear 96-well plates in Tecan Spark or Sunrise plate readers, incubating at 37°C with shaking and measuring A_600_ at 30-minute intervals for 24 hours.

**TABLE 1.** Strains and plasmids used in this study. Unless otherwise indicated, strains and plasmids were generated in the course of this work. Abbreviations: Ap^R^, ampicillin resistance; Cm^R^, chloramphenicol resistance; Kn^R^, kanamycin resistance; Rif^R^, rifampicin resistance.

| Strain | Marker(s) | Relevant Genotype | Source |
| --- | --- | --- | --- |
| <u>E. coli strains:</u> |  |  |  |
| MG1655 |  | F<sup>-</sup>, λ<sup>-</sup>, *rph-1 ilvG<sup>-</sup> rfb-50* | (115) |
| CF1693(M+) | Cm<sup>R</sup> Kn<sup>R</sup> | MG1655 λ<sup>+</sup> φ80<sup>+</sup> *relA251::kan<sup>+</sup> spoT207::cat<sup>+</sup>*<br>*rpoB1693* (encoding RpoB<sup>N1129K</sup>) *rpoD1693*<br>(encoding RpoD<sup>D64G</sup>) | (116, 117) |
| BH330 | Ap<sup>R</sup> | MG1655 λ*attB::P<sub>lac</sub>-gfp-ftsZ*, *bla<sup>+</sup>* | (84) |
| MJG0224 |  | MG1655 Δ*ppk-749* | (40) |
| MJG0226 |  | MG1655 Δ*relA782* | (42) |
| MJG0344 |  | MG1655 Δ*rpoS746* | (42) |
| MJG1090 | Kn<sup>R</sup> | MG1655 Δ*ppk-749* Δ*relA782::kan<sup>+</sup>* |  |
| MJG1097 | Kn<sup>R</sup> | MG1655 Δ*rpoS746* Δ*relA782::kan<sup>+</sup>* |  |
| MJG1116 |  | MG1655 Δ*ppk-749* Δ*relA782* |  |
| MJG1119 |  | MG1655 Δ*rpoS746* Δ*relA782* |  |
| MJG1136 | Cm<sup>R</sup> | MG1655 $\Delta relA782 spoT207::cat^+$ | (42) |
| MJG1137 | Cm<sup>R</sup> | MG1655 $\Delta ppk-749 \Delta relA782 spoT207::cat^+$ | |
| MJG1236 | Rif<sup>R</sup> | MG1655 $\Delta ppk-749 \Delta relA782 rpoB3449$<br>(encoding RpoB<sup>$\Delta Ala532$</sup>) | |
| MJG1237 | Cm<sup>R</sup> Rif<sup>R</sup> | MG1655 $\Delta relA782 spoT207::cat^+ rpoB3449$<br>(encoding RpoB<sup>$\Delta Ala532$</sup>) | (42) |
| MJG1241 | Cm<sup>R</sup> Rif<sup>R</sup> | MG1655 $\Delta ppk-749 \Delta relA782 spoT207::cat^+$<br>$rpoB3449$ (encoding RpoB<sup>$\Delta Ala532$</sup>) | |
| MJG1282 | Kn<sup>R</sup> | MG1655 $\Delta ppk-749 \Delta relA782 \Delta spoT1000::kan^+$ | |
| MJG1287 | Kn<sup>R</sup> | MG1655 $\Delta relA782 \Delta spoT1000::kan^+$ | (42) |
| MJG1575 | Rif<sup>R</sup> | MG1655 $\Delta relA782 rpoB3443$ (encoding<br>RpoB<sup>L533P</sup>) | |
| MJG1576 | Rif<sup>R</sup> | MG1655 $\Delta relA782 rpoB148$ (encoding<br>RpoB<sup>D517V</sup>) | |
| MJG1577 | Rif<sup>R</sup> | MG1655 $\Delta ppk-749 \Delta relA782 rpoB3443$<br>(encoding RpoB<sup>L533P</sup>) | |
| MJG1578 | Rif<sup>R</sup> | MG1655 $\Delta ppk-749 \Delta relA782 rpoB148$<br>(encoding RpoB<sup>D517V</sup>) | |
| MJG1579 | Cm<sup>R</sup> Rif<sup>R</sup> | MG1655 $\Delta relA782 spoT207::cat^+ rpoB3443$<br>(encoding RpoB<sup>L533P</sup>) | |
| MJG1580 | Cm<sup>R</sup> Rif<sup>R</sup> | MG1655 $\Delta relA782 spoT207::cat^+ rpoB148$<br>(encoding RpoB<sup>D517V</sup>) | |
| MJG1581 | Cm<sup>R</sup> Rif<sup>R</sup> | MG1655 $\Delta ppk-749 \Delta relA782 spoT207::cat^+$ | |
|  |  | *rpoB3443* (encoding RpoB<sup>L533P</sup>) |  |
| MJG1582 | Cm<sup>R</sup> Rif<sup>R</sup> | MG1655 $\Delta ppk-749 \Delta relA782 spoT207::cat^+$<br>*rpoB148* (encoding RpoB<sup>D517V</sup>) | |
| MJG2401 | Ap<sup>R</sup> | MG1655 $\lambda attB::P_{lac}-gfp-ftsZ, bla^+$ | |
| MJG2402 | Ap<sup>R</sup> | MG1655 $\Delta ppk-749 \lambda attB::P_{lac}-gfp-ftsZ, bla^+$ | |
| MJG2403 | Kn<sup>R</sup> Ap<sup>R</sup> | MG1655 $\Delta ppk-749 \Delta relA782::kan^+ \lambda attB::P_{lac}-gfp-ftsZ, bla^+$ | |
| MJG2404 | Kn<sup>R</sup> Ap<sup>R</sup> | MG1655 $\Delta relA782 \Delta spoT1000::kan^+ \lambda attB::P_{lac}-gfp-ftsZ, bla^+$ | |
| MJG2405 | Kn<sup>R</sup> Ap<sup>R</sup> | MG1655 $\Delta ppk-749 \Delta relA782 \Delta spoT1000::kan^+ \lambda attB::P_{lac}-gfp-ftsZ, bla^+$ | |
| **Plasmid** | **Marker(s)** | **Relevant Genotype** | **Source** |
| pUC18 | Ap<sup>R</sup> | *bla*<sup>+</sup> | (126) |
| pKD46 | Ap<sup>R</sup> | $\lambda$ Red<sup>+</sup>, *bla*<sup>+</sup> | (121) |
| pKD4 | Kn<sup>R</sup> | *kan*<sup>+</sup> | (121) |
| pCP20 | Ap<sup>R</sup> Cm<sup>R</sup> | Flp<sup>+</sup> *bla*<sup>+</sup> *cat*<sup>+</sup> | (121) |
| pPPK1 | Cm<sup>R</sup> | *ppk*<sup>+</sup> *cat*<sup>+</sup> | (40) |
| pPPK8 | Ap<sup>R</sup> | *ppk*<sup>+</sup> *bla*<sup>+</sup> |  |
| pSPOT1 | Ap<sup>R</sup> | *spoT*<sup>+</sup> *bla*<sup>+</sup> |  |
| pSPOT2 | Ap<sup>R</sup> | *spoT*<sup>G217A, C219T</sup> (encoding SpoT<sup>D73N</sup>) *bla*<sup>+</sup> |  |
| pSPOT3 | Ap<sup>R</sup> | *spoT*<sup>G775A, C777T</sup> (encoding SpoT<sup>D259N</sup>) *bla*<sup>+</sup> |  |
| pSPOT4 | Ap<sup>R</sup> | *spoT*<sup>G217A, C219T, G775A, C777T</sup> (encoding SpoT<sup>D73N</sup>,<br>D259N) |  |
|  |  | D259N)<br>*bla*<sup>+</sup> |  |

### Strain construction

All *E. coli* strains used in this study were derivatives of wild-type strain MG1655 (F^-^, λ^-^, *rph-1 ilvG^-^ rfb-50*) (115). We confirmed chromosomal mutations by PCR and whole-genome sequencing (SeqCenter, Philadelphia, PA). All strains derived from CF1693(M+)(116, 117) were confirmed free of contamination with λ and ϕ80 phage by PCR (42).

We used P1*vir* phage transduction (118, 119) to move the Δ*relA782*::*kan*^+^ allele from the Keio collection (120) into strains MJG0224 (Δ*ppk*-*749*)(40) and MJG0344 (Δ*rpoS746*)(42) to generate strains MJG1090 (Δ*ppk*-*749* Δ*relA782*::*kan*^+^) and MJG1097 (Δ*rpoS746* Δ*relA782*::*kan*^+^). We then used plasmid pCP20 (121) to resolve the kanamycin resistance cassettes in those strains, generating strains MJG1116 (Δ*ppk*-*749* Δ*relA782*) and MJG1119 (Δ*rpoS746* Δ*relA782*). We used P1*vir* phage transduction to move the *spoT207*::*cat*^+^ allele from CF1693(M+)(116, 117) into MJG1116 (Δ*ppk*-*749* Δ*relA782*), generating strain MJG1137 (Δ*ppk*-*749* Δ*relA782 spoT207*::*cat*^+^). The *spoT* gene of strain MJG1116 (Δ*ppk*-*749* Δ*relA782*) was replaced with a pKD4-derived kanamycin resistance cassette by recombineering (121) using primers 5’ GTT ACC GCT ATT GCT GAA GGT CGT CGT TAA TCA CAA AGC GGG TCG CCC TTG GTG TAG GCT GGA GCT GCT TC 3’ and 5’ GGC GAG CAT TTC GCA GAT GCG TGC ATA ACG TGT TGG GTT CAT AAA ACA TTA CAT ATG AAT ATC CTC CTT AG 3’, yielding strain MJG1282 (Δ*ppk*-*749* Δ*relA782 spoT1000*::*kan*^+^).

Oligo-directed recombineering (122) was used to construct chromosomal *rpoB3449*, *rpoB3443*, and *rpoB148* alleles (42, 69, 91, 123–125) using the mutagenic primers 5’ CCT GCA CGT TCA CGG GTC AGA CCG CCT GGA CCA AGA GAA ATA CGA CGT TTG TGC GTA ATC TCA GAC AGC G 3’, 5’ AAG CCT GCA CGT TCA CGG GTC AGA CCG CCG GGA CCG GGG GCA GAG ATA CGA CGT TTG TGC GTA ATC TCA GAC A 3’, and 5’ ATA CGA CGT TTG TGC GTA ATC TCA GAC AGC GGA TTA TTT TGG ACC ATA AAC TGA GAC AGC TGG CTG GAA CCG A 3’ respectively, each of which contained four 5’ phosphorothiorate linkages to stabilize the primers. The *rpoB3449* primer deletes nucleotides G1593 through A1596 of *rpoB*, removing the codon for alanine 532 of RpoB, and also incorporates silent mutations in four adjacent codons (C1590T, C1593T, C1599T, and C1602T) to avoid mismatch repair (42). The *rpoB3443* primer mutates nucleotide T1598 of *rpoB* to C, changing leucine 533 to proline, and incorporates silent mutations in four adjacent codons (C1593T, A1596C, C1602T, and A1605C). The *rpoB148* primer mutates nucleotide A1547 of *rpoB* to T, changing aspartic acid 516 to valine, and incorporates silent mutations in three adjacent codons (G1551A, C1554T, and C1557T). Strains MJG0226 (Δ*relA782*) or MJG1116 (Δ*ppk*-*749* Δ*relA782*) were transformed with pKD46 (121), induced to express the λ Red recombinase, and electroporated with 250 pmol of mutagenic primer. Recombinant colonies were selected at 37°C on LB plates containing rifampicin. The sequence of *rpoB* alleles was confirmed by PCR amplification of a fragment of *rpoB* with primers 5’ GAT GTT ATG AAA AAG CTC 3’ and 5’ CTG GGT GGA TAC GTC CAT 3’ and Sanger sequencing of the resulting product (UAB Heflin Sequencing Core Facility). After curing pKD46 by growth at 37°C, this yielded strains MJG1236 (Δ*ppk-749* Δ*relA782 rpoB3449*), MJG1575 (Δ*relA782 rpoB3443*), MJG1576 (Δ*relA782 rpoB148*), MJG1577 (Δ*ppk-749* Δ*relA782 rpoB3443*), and MJG1578 (Δ*ppk-749* Δ*relA782 rpoB148*).

We used P1*vir* phage transduction to move the *spoT207*::*cat*^+^ allele from CF1693(M+)(116, 117) into MJG1236 (Δ*ppk-749* Δ*relA782 rpoB3449*), generating strain MJG1237 (Δ*ppk-749* Δ*relA782 spoT207*::*cat*^+^ *rpoB3449*). We used P1*vir* phage transduction to move the *spoT207*::*cat*^+^ allele from MJG1136 (Δ*relA782 spoT207*::*cat*^+^)(42) into strains MJG1575 (Δ*relA782 rpoB3443*), MJG1576 (Δ*relA782 rpoB148*), MJG1577 (Δ*ppk-749* Δ*relA782 rpoB3443*), and MJG1578 (Δ*ppk-749* Δ*relA782 rpoB148*), yielding strains MJG1579 (Δ*relA782 spoT207*::*cat*^+^ *rpoB3443*), MJG1580 (*relA782 spoT207*::*cat*^+^ *rpoB148*), MJG1581 (Δ*ppk-749* Δ*relA782 spoT207*::*cat*^+^ *rpoB3443*), and MJG1582 (Δ*ppk-749* Δ*relA782 spoT207*::*cat*^+^ *rpoB148*) respectively.

To construct fluorescent reporter strains, we used P1*vir* phage transduction to move the λ*attB::P_lac_-gfp-ftsZ* allele from strain BH330 (84) into strains MG1655, MJG0224 (Δ*ppk-749*), MJG1090 (Δ*ppk-749* Δ*relA782*::*kan*^+^), MJG1287 (Δ*relA782* Δ*spoT1000*::*kan*^+^), or MJG1282 (Δ*ppk-749* Δ*relA782* Δ*spoT1000*::*kan*^+^), selecting for the linked ampicillin resistance marker. This resulted in strains MJG2401 (λ*attB::P_lac_-gfp-ftsZ*, *bla*^+^), MJG2402 (Δ*ppk-749* λ*attB::P_lac_-gfp-ftsZ*, *bla*^+^), MJG2403 (Δ*ppk-749* Δ*relA782*::*kan*^+^ λ*attB::P_lac_-gfp-ftsZ*, *bla*^+^), MJG2404 (Δ*relA782* Δ*spoT1000*::*kan*^+^ λ*attB::P_lac_-gfp-ftsZ*, *bla*^+^), and MJG2405 (Δ*ppk-749* Δ*relA782* Δ*spoT1000*::*kan*^+^ λ*attB::P_lac_-gfp-ftsZ*, *bla*^+^).

### Plasmid construction

The *E. coli* MG1655 *ppk* coding sequence (2,067 bp) plus 20 bp of upstream sequence was subcloned from pPPK1 (40) into the *Kpn*I and *Hin*dIII sites of plasmid pUC18 (126) to generate plasmid pPPK8 (*ppk*^+^ *bla*^+^). The *spoT* coding sequence (2,109 bp) was amplified from *E. coli* MG1655 genomic DNA with primers 5’ AGA TCT AGA TTG TAT CTG TTT GAA AGC CTG AAT C 3’ and 5’ CTT AAG CTT TTA ATT TCG GTT TCG GGT GAC 3’ and cloned into the *Xba*I and *Hin*dIII sites of plasmid pUC18 (126) to generate plasmid pSPOT1 (*spoT*^+^ *bla*^+^). We used single primer site-directed mutagenesis (127) to mutate pSPOT1 with primers 5’ GGC GGC GCT GCT GCA TAA TGT GAT TGA AGA TAC TCC 3’ and 5’ CGT TTT CAC TCG ATC ATG AAT ATC TAC GCT TTC CGC GTG 3’. This yielded pSPOT2, containing a *spoT*^G217A,C219T^ allele (encoding SpoT^D73N^), and pSPOT3, containing a *spoT*^G775A,C777T^ allele (encoding SpoT^D259N^). We then used single primer site-directed mutagenesis (127) to further mutate pSPOT3 with primer 5’ GGC GGC GCT GCT GCA TAA TGT GAT TGA AGA TAC TCC 3’. This yielded pSPOT4, containing a *spoT*^G217A,C219T,G775A,C777T^ allele (encoding SpoT^D73N, D259N^)

### (p)ppGpp Quantification

Cultures were grown overnight in LB, then sub-cultured into 5 mL of LB for each sample, and grown with shaking at 37°C for 2-3 hours until cells started to grow exponentially (OD_600_ = ∼ 0.1). Cells were put into two different 50 mL conical tubes containing 2 mL of the strain of interest, rinsed three times with sterile PBS, and resuspended in MOPS glucose medium. To one tube we added add ^32^P (Phosphorus-32 Radionuclide, 1mCi (37 MBq) ®Revvity) to a final concentration of 20 µCi/mL (128). Cultures were incubated with shaking at 37 °C until they reached an OD_600_ between 0.4-0.5 (for strains able to replicate in MOPS medium), then 200 µL culture was added to 40 µL of 2 M formic acid (129), incubated on ice for 15-60 min, then centrifuged at 16,000 x g for 2 minutes at 4°C and supernatants were stored at -20°C. Thin-layer chromatography (TLC) was carried out to visualize and quantify phosphorylated guanine nucleotides as previously described (129, 130). TLC plates were analyzed on a Typhoon Biomolecular Imager, Cytiva. Spot intensity was quantified using ImageQuant, and (p)ppGpp quotient was expressed as a fraction of the total guanosine pool, i.e. (pppGpp+ppGpp)/(pppGpp+ppGpp+GTP+GDP) (130).

### Size fractionation of yeast extract

We dissolved yeast extract in water (0.25 g ml^-1^) to make a concentrated stock solution. We placed the resulting solution in a 3500 Da MWCO Slide-A-Lyzer cassette (ThermoFisher) and dialyzed it against distilled water to remove components smaller than 3500 Da. Similarly, we used a 3000 Da MWCO Amicon™ Ultra-15 Centrifugal Filter (Millipore Sigma) unit to remove yeast extract stock solution components larger than 3000 Da. The resulting solutions were filter-sterilized and used to supplement minimal media at the indicated concentrations.

### Fluorescent Time-Lapse Microscopy

Strains of interest were grown overnight in 5 mL of LB liquid broth shaking at 37°C to obtain a saturated culture. Cultures were back diluted 1,000-fold and grown to early exponential phase (approximately 3 hours). Cells were concentrated by centrifugation, when necessary, at 5,000 *g* for 1 minute, before being resuspended in 100 µL of media. Cells were immobilized on agarose pads by spotting 0.5 µL of concentrated culture on pad, then inverting the pad onto a glass bottom petri dish for imaging. Agarose pads for microscopy were constructed out of LB containing 1.5% agarose (Thermo Fisher cat. #16500500). Imaging was performed using equipment available at the University of Alabama at Birmingham High-Resolution Imaging Facility (HRIF); a Nikon Ti2 inverted fluorescence microscope with a tandem galvano and Nikon A1R-HD25 resonance scanner up to 30 1024x1024 images/sec.

Tokai Hit incubation stage chamber was used to heat samples and objectives to 37°C to facilitate growth of bacteria for live imaging. Images were captured at 60x or 100x magnification in both transmitted light differential interference contrast image and GFP or appropriate fluorescent channels as needed. Automated time lapse imaging was performed at 37°C, and motorized x, y, and z tracking was controlled and automated by acquisition software Nis Elements 5.0 Imaging Software available in the HRIF at UAB. Analysis of microscopy images captured will be analyzed in FIJI (**F**iji **I**s **J**ust **I**mageJ) for cell length, fluorescence quantification and tracking (131–133).

1mM Isopropyl β-d-1-thiogalactopyranoside (IPTG) was added to cells containing the λ*attB::P_lac_-gfp-ftsZ* reporter. 1 mM IPTG was also added to the agarose pad for imaging during the time-lapse of these strains.

The fraction of cells that developed into branching cells was determined from observing time-lapse microscopy. Cells were grown on LB agarose pads at 37°C for 3 hours, and when branched cells formed, time-lapse was stopped and the total number of cells was counted, and the number of cells that were branched.

### Cryo-Electron Microscopy

Cryo-Electron Microscopy was performed with the help of the UAB institutional Research Core Program with the help of Dr. Terje Dokland and Dr. James Kizziah (University of Alabama at Birmingham). A Thermo Fisher Scientific (TFS) Glacios 2 equipped with a Falcon 4i direct electron detector is optimized for high-throughput cryo-EM and ease-of-use via the TFS EPU and Tomography software. It has demonstrated data collection speeds up to ∼500 images/hr and capability of 2.2Å resolution via single particle analysis with apoferritin. A TFS Talos F200C equipped with a Ceta-S CMOS camera and a Direct Electron Apollo direct electron detector is used for imaging of negatively stained samples and specialized cryo-EM applications.

For sample preparation, a Pelco EasiGlow glow discharge machine, a PIE Scientific TergeoEM plasma cleaner, and FEI Vitrobot Mark IV sample vitrification robot were used. Gatan 626 and 698 “Elsa” cryo-holders are used for cryo-EM on the Talos F200C. An in-house GPU-accelerated computing workstation is used for on-the-fly processing of single particle cryo-EM data with CryoSPARC Live, and the CEMF has a direct 10Gbps fiber link to the UAB supercomputer for offloading and distribution of data to users. Cells were grown overnight until the morning when they were back diluted and grown for 3 hours until an OD_600_ of 0.1 was reached, and cells were centrifuged at 8,000 g for 2 minutes, and resuspended in PBS for CEM sample prep.

### Transmission Electron Microscopy (TEM)

TEM was performed at/by the High-Resolution Imaging Facility (HRIF) at the University of Alabama at Birmingham. Wild type MG1655 (MJG0001) and *ppk relA spoT* (MJG1282) were grown in LB at 37°C until exponential phase growth (approximately OD_600_ 0.3 – 0.5). Cells were then spun down and collected. Remove media from pellet and fix in 1% Osmium tetroxide (EMS) in 0.1M Sodium Cacodylate Buffer pH 7.4 at room temperature in the dark for 1 hour, then 3 times 0.1M Sodium Cacodylate Buffer pH 7.4 rinse for 15 minutes each. 1% Low molecular weight tannic acid (Ted Pella Inc) for 20 minutes, 3 times 0.1M Sodium Cacodylate Buffer pH 7.4 rinse for 15 minutes each.

The specimens are dehydrated through a series of graded ethyl alcohols from 50 to 100%. The schedule is as follows: 50% for 5 min., 2% uranyl acetate in 50% EtOH for 30 minutes in dark, 50% for 5 min, 80% for 5 min, 95% for 5 min, and four changes of 100% for 15 minutes each. After dehydration the infiltration process requires steps through an intermediate solvent, 2 changes of 100% propylene oxide (P.O.) for 10 minutes each and finally into a 50:50 mixture of P.O. and the embedding resin (Embed 812, Electron Microscopy Sciences, Hatfield, PA) for 12-18 hours.

The specimen is transferred to fresh 100% embedding media. The following day the specimen is then embedded in a fresh change of 100% embedding media. Blocks polymerize overnight in a 60 degree C embedding oven and are then ready to section.

### Procedure to Section for Transmission Electron Microscopy

The resin blocks are first thick sectioned at 0.5-1 microns with a diamond histo knife using an ultramicrotome and sections are stained with Toluidine Blue, these sections are used as a reference to trim blocks for thin sectioning. The appropriate blocks are then thin sectioned using a diamond knife (Diatome, Electron Microscopy Sciences, Fort Washington, PA)) at 70-100nm (silver to pale gold using color interference) and sections are then placed on either copper or nickel mesh grids. After drying, the sections are stained with heavy metals, uranyl acetate and lead citrate for contrast. After drying the grids are then viewed on a JEOL 1400 FLASH 120kv TEM (JEOL USA Inc, Peabody, MA). Digital images are taken with an AMT NanoSprint43 Mark II camera (AMT Imaging, Woburn, MA) and transferred via UAB BOX or other device.

### Statistical analyses

We used GraphPad Prism version 10.2.2 for Macintosh (GraphPad Software) to perform all statistical analyses and graph generation.

### Data availability

All strains generated in the course of this work are available from the authors upon request. We deposited DNA sequencing data in the NIH Sequence Read Archive (accession number: PRJNA1032912), and all other raw data is available on FigShare (DOI: 10.6084/m9.figshare.c.7430740).

## ACKNOWLEDGEMENTS

This project was supported by University of Alabama at Birmingham Department of Microbiology development funds and by NIH grants R35 GM124590 (to MJG) and K12 GM088010 (to CWH). The authors have no conflicts of interest to declare. We would like to thank Drs. Petra Levin and Sarah Anderson (Washington U.) for the gift of fluorescent reporter strains and for helpful discussions and Dr. David Schneider (UAB Department of Biochemistry and Molecular Genetics) for assistance with (p)ppGpp measurements.

